# A binuclear copper enzyme platform for enantioconvergent radical (pseudo)halogenation

**DOI:** 10.64898/2026.09.20.753013

**Authors:** Xiao-Wang Chen, Oluwatosin Maryam Adeyemo, Heyu Chen, Hongda Chen, Ngoc Nguyen, Joseph Palazzo, Liu-Peng Zhao, Ken Lin, Binh Khanh Mai, Peng Liu, Shiliang Tian, Yang Yang

## Abstract

Despite their intriguing native metalloenzymology, naturally occurring copper enzymes remain largely underexploited for new-to-nature biocatalytic reactions. Herein, we report the systematic investigation and reprogramming of natural copper enzymes to catalyze unnatural free radical (pseudo)halogenation reaction in a highly enantioselective fashion. Evaluating Cu enzymes in the decarboxylative azidation of redox-active esters revealed activity across multiple Cu enzyme families, with type III binuclear Cu enzymes, particularly the *Bacillus megaterium* tyrosinase (*Bm*Tyr), exhibiting superior activity and enantioselectivity across both stabilized and unstabilized alkyl radicals upon further engineering. The strong halide binding affinity of the binuclear Cu system also enabled challenging enantioconvergent bromination, chlorination and isothiocyanation reactions, which remained inaccessible to repurposed nonheme Fe enzymes. Further EPR and UV– visible spectroscopic analyses confirmed the coupled binuclear nature of wild-type and engineered bacterial tyrosinases and provided insights into the origin of their enhanced activity. Collectively, this study establishes binuclear copper enzymes as a powerful platform for new-to-nature stereoselective radical reactions, expanding the scope of metalloenzyme catalysis beyond mononuclear systems.

## Introduction

Over the past decade, metalloenzymes have been reprogrammed and evolved into powerful biocatalysts for synthetically valuable stereoselective transformations that operate through mechanisms unknown in nature^1-7^. In particular, recent work from our laboratory and other researchers has established both heme^8-14^ and nonheme^15-27^ Fe enzymes as tunable platforms for a wide range of stereoselective radical transformations, providing a new solution to the longstanding challenge of imposing stereocontrol over free radical intermediates. In contrast to the extensive development of Fe-based systems, naturally occurring metalloenzymes that utilize alternative first-row transition-metal cofactors, particularly copper^28^, remain largely underexploited for non-native radical biocatalysis^7^. Although metal substitution in nonheme Fe and Zn enzymes has enabled access to exciting new reactivities^29-34^, the native coordination environments of these proteins are evolutionarily optimized for Fe or Zn and may not provide an ideal environment for alternative metal ions such as Cu, thereby posing a key limitation of this strategy.

To address these limitations, we set out to systematically evaluate the synthetic potential of natural copper enzymes for enabling new-to-nature radical reactions via a metalloredox mechanism, particularly those that remain challenging to access using reprogrammed Fe enzymes (Fig. 1). In nature, a diverse array of type I, II and III copper enzymes, featuring distinct coordination chemistry (Fig. 1a), have evolved to catalyze various oxidative transformations using O_2_ or H_2_O_2_ as the terminal oxidant^28^. In contrast to native enzymology, we sought to investigate the potential of Cu enzymes to promote overall redox-neutral transformations under anaerobic conditions via Cu(II)/Cu(I) redox cycling. In particular, we were intrigued by the possibility that copper enzymes could catalyze radical (pseudo)halogenation reactions, including chlorination, bromination, azidation, isothiocyanation and thiocyanation, of non-native substrate classes via a radical rebound mechanism. For several decades, the metallohalogenase literature has been dominated by α-ketoglutarate-dependent nonheme Fe enzymes^35,36^. In 2025, groundbreaking work by Tang and coworkers reported a rare example of Cu-dependent biosynthetic enzyme capable of catalyzing C–H chlorination of atpenin B (Fig. 1b)^37^, ushering in a new frontier of metallohalogenase research. From a chemical reactivity perspective, inspired by the enhanced (pseudo)halide binding affinity of Cu(II) centers arising from their softer character relative to Fe(III), we envisioned that copper enzymes could enable a broader range of challenging (pseudo)halogenation reaction.

**Fig. 1.**
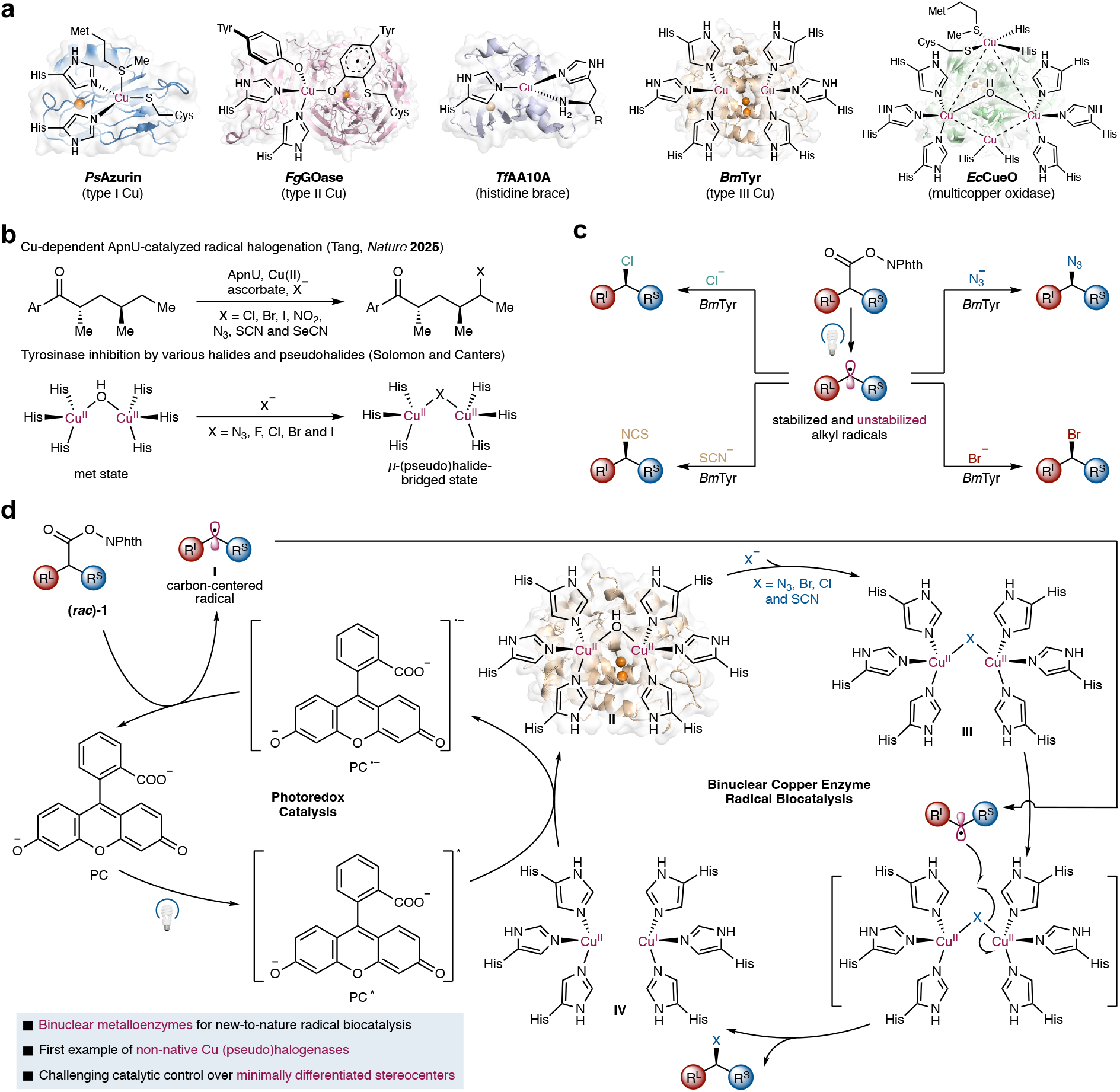
Binuclear copper enzymes as a powerful platform for advancing new-to-nature stereoselective radical reactions. **a**, Coordination chemistry in mono- and binuclear Cu enzymes. **b**, Natural Cu-dependent halogenase and tyrosinase inhibition by halides. **c**, New-to-nature biocatalytic enantioconvergent radical (pseudo)halogenation reactions, including azidation, bromination, chlorination and isothiocyanation. **d**, Proposed mechanism of biocatalytic radical (pseudo)halogenation reactions using binuclear copper enzymes. R^L^: larger substituent, R^S^: smaller substituent.

Among natural copper enzymes, we were particularly fascinated by the largely underexplored synthetic potential of binuclear copper enzymes. In native enzymology, bimetallic active sites synergistically harness the redox properties of two proximal metal centers to enable multielectron transformations through mechanisms inaccessible to mononuclear metalloenzymes^38^. To date, such bimetallic systems, including diiron^39^ and dicopper enzymes^40^, have rarely been leveraged for new-to-nature biocatalysis, limiting the exploration of their multimetallic cooperativity for challenging reactivity. In particular, important studies by Solomon^41^ and Canters^42,43^ revealed that tyrosinases, members of the type III copper enzyme superfamily, bind (pseudo)halide ions cooperatively to form μ-(pseudo)halide-bridged dicopper intermediates (Cu–X–Cu, X = N_3_, F, Cl, Br and I) (Fig. 1b). Although (pseudo)halide binding is typically inhibitory to native tyrosinase activity, we envisioned that this strong binding affinity, combined with the unique reactivity of μ-(pseudo)halide-bridged dicopper intermediates, could be harnessed strategically to enable challenging enantioselective radical (pseudo)halogenation chemistry^44,45^. In particular, by leveraging the exquisite enzymatic stereocontrol over transient free radical intermediates, this dicopper enzyme platform holds the potential to construct minimally differentiated alkyl–alkyl stereocenters, which continue to present substantial synthetic hurdles for state-of-the-art chiral small-molecule copper catalysts^46,47^.

Our envisioned dual photobiocatalytic cycles for dicopper enzyme-catalyzed radical (pseudo)halogenation are described in Fig. 1d. In the photoredox cycle, the use of an appropriate photosensitizer would generate an alkyl radical intermediate (**I**) from the corresponding *N*-hydroxyphthalimide ester substrate **1**. Concurrently, in the biocatalytic cycle, the dicopper enzyme (**II**) binds (pseudo)halide ions, including challenging halides such as bromide and chloride, to form a μ-(pseudo)halide-bridged Cu(II)_2_ intermediate (**III**). The photoredox-generated alkyl radical (**I**) would enter the enzyme active site and react with the copper-bound bridging μ-(pseudo)halide, thereby forming the radical (pseudo)halogenation products (**2**–**5**) and generating a mixed-valence Cu(II)/Cu(I) dicopper enzyme state (**IV**). We hypothesized that electronic coupling between the two proximal Cu(II)/Cu(I) centers would stabilize this mixed-valence intermediate (**IV**) ^28,48,49^, thereby facilitating otherwise challenging radical halogenation reactions. Finally, single-electron oxidation of the Cu(II)/Cu(I) state enzyme (**IV**) by the photoredox catalyst would regenerate the dicopper enzyme (**II**), thereby completing the catalytic cycle. Herein, we report the successful implementation of this design through the development of dicopper enzyme-catalyzed enantioconvergent (pseudo)halogenation reactions. This platform enables a diverse set of asymmetric radical transformations, including bromination, chlorination and isothiocyanation, which have remained challenging for chiral small-molecule copper catalysts (Fig. 1c). These dicopper enzyme-based transformations expand the chemical landscape of new-to-nature nonheme Fe-based biocatalysis^19,20^, while providing mechanistic insights into recently discovered copper-dependent biosynthetic halogenation pathways^37^.

## Results and discussion

### Evaluation of natural copper enzymes for asymmetric radical azidation and discovery of binuclear Cu tyrosinases as highly effective biocatalysts

To evaluate the potential of naturally occurring Cu-dependent enzymes^28^ in promoting unnatural radical (pseudo)halogenation reactions, we built an in-house collection of various type I, II and III copper proteins and multinuclear copper oxidases (MCOs)^28^ that could be conveniently expressed in *Escherichia coli* at high expression levels (Fig. 2). This Cu enzyme library encompassed type I blue copper proteins including azurin, amicyanin and their mutants; type II copper enzymes including galactose oxidases (GOases) and copper amine oxidases (CAOs); type III copper enzymes including various tyrosinases; as well as MCOs, including laccases. Collectively, our in-house Cu-dependent protein library includes mononuclear, binuclear and multinuclear Cu enzymes with diverse Cu coordination chemistry, thereby setting the stage for the discovery of new-to-nature Cu enzyme-catalyzed reactivity.

**Fig. 2.**
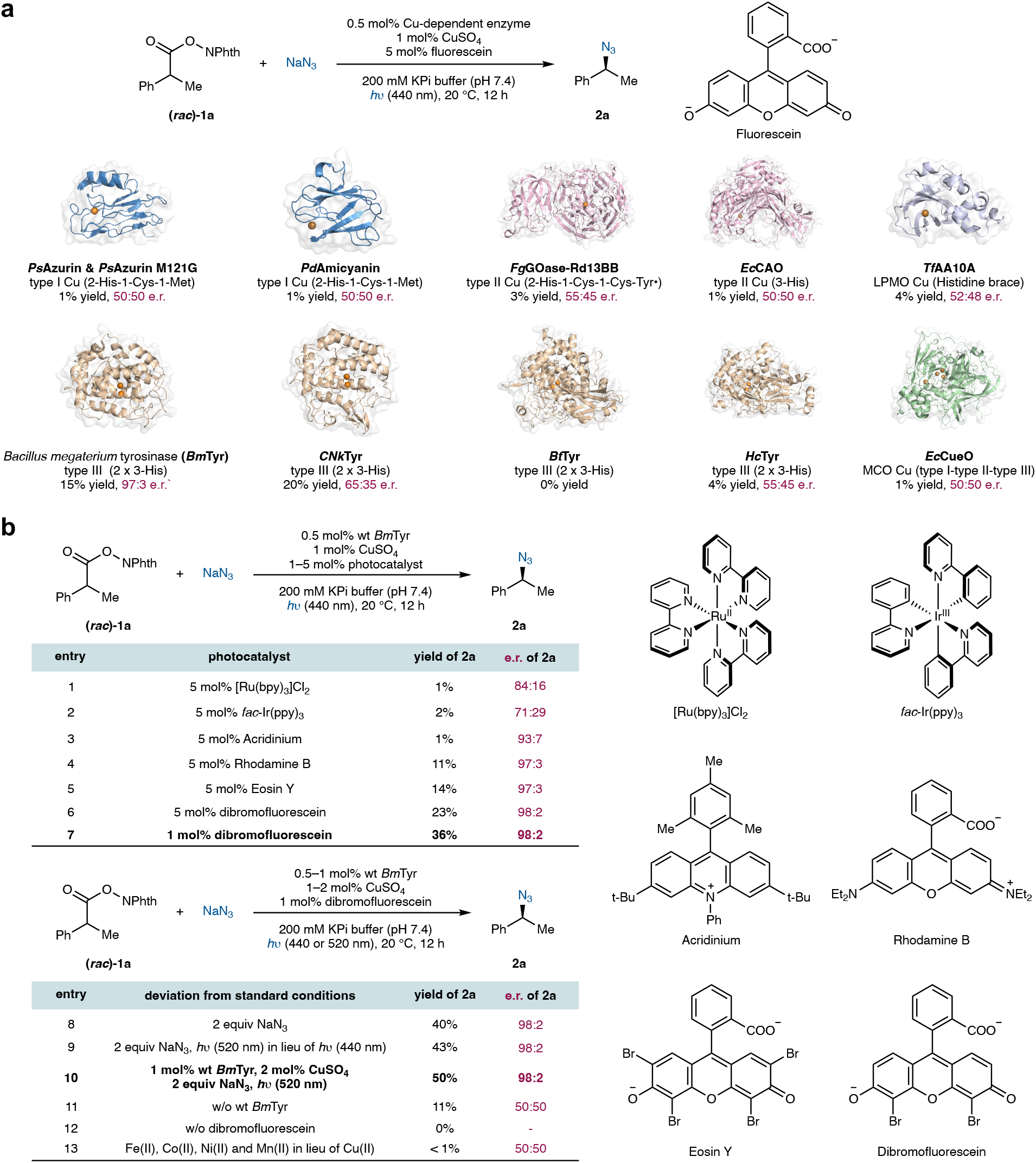
Discovery and optimization of copper-dependent enzyme-catalyzed enantioselective photometallobiocatalytic decarboxylative azidation. **a**, Evaluation of natural Cu enzymes for photometallobiocatalytic azidation. **b**, Optimization of photocatalysts and control experiments. Reaction conditions: **1a** (1.0 equiv, 4.0 mM), NaN_3_ (2.0–5.0 equiv, 4.0–20.0 mM), 0.5–1 mol% Cu-dependent enzyme (20–40 μM), 1 mol% CuSO_4_ (40 μM), 1–5 mol% photocatalyst (40 μM– 200 μM), *h*υ (440 or 520 nm), KPi buffer (200 mM, pH 7.4), 20 °C, 12 h.

To investigate whether naturally occurring Cu enzymes could facilitate radical-mediated (pseudo)halogenation reactions, we first evaluated their catalytic activity in photobiocatalytic azidation using racemic *N*-hydroxyphthalimide ester (*rac*)-**1a** as the radical precursor. Under visible light irradiation (440 nm), in the presence of 5 mol% fluorescein as the photosensitizer, the use of type I blue copper proteins, including azurin^50^ from *Pseudomonas aeruginosa* and amicyanin^51^ from *Paracoccus denitrificans* as well as their single mutant azurin M121G^52^ possessing an open coordination site, provided <1% yield of **2a** with no enantioselectivity. The type II copper enzyme *Escherichia coli* copper amine oxidase (*Ec*CAO)^53^ also showed minimal azidation activity. Interestingly, other type II copper enzymes exhibited low levels of activity and measurable enantioselectivity. The use of a thermophilic lytic polysaccharide monooxygenase from *Thermobifida fusca* (*Tf*AA10A)^54^ afforded **2a** in 4% yield and 52:48 e.r., while the employment of an engineered variant of galactose oxidase from *Fusarium graminearum* (*Fg*GOase)^55^ gave rise to **2a** in 3% yield and 55:45 e.r.. Collectively, these results indicated the potential of these type II mononuclear Cu enzymes in promoting asymmetric radical azidation.

Among all the natural Cu enzymes we evaluated, bacterial tyrosinases, a class of type III binuclear Cu enzymes featuring a coupled binuclear Cu center, displayed the highest initial activity and enantioselectivity. In particular, tyrosinase from *Bacillus megaterium* (*Bm*Tyr)^56^ furnished azidation product **2a** in 15% yield and excellent enantioselectivity (97:3 e.r.), clearly demonstrating enzymatic conversion of (*rac*)-**1a**. The archaeal tyrosinase from *Candidatus Nitrosopumilus koreensis* (*CNk*Tyr)^57^ possessing a more solvent exposed active site exhibited enhanced activity, providing **2a** in 20% yield albeit with a slightly reduced enantioselectivity (65:35 e.r.). Other tyrosinases, including the *Burkholderia thailandensis* tyrosinase (*Bt*Tyr)^58^ and *Hahella* sp. CCB MM4 tyrosinase (*Hc*Tyr)^59^ afforded **2a** in 0% yield and 4% yield (55:45 e.r.), respectively. Given the excellent stereocontrol and initial activity of *Bm*Tyr, this type III binuclear Cu enzyme was selected for further optimization and protein engineering.

Using 0.5 mol% *Bm*Tyr as the biocatalyst, we next evaluated a range of photocatalysts in the current photobiocatalytic asymmetric azidation. Transition metal-based photocatalysts, showed low levels of activity and reduced enantioselectivity (Fig. 2b, entries 1 and 2). Although the use of an acridinium salt also afforded low yields, excellent enantioselectivity of **2a** was observed (93:7 e.r., entry 3). Organic photocatalysts from the rhodamine family such as rhodamine B delivered **2a** in 11% yield and 97:3 e.r. (entry 4). Other organic dyes from the fluorescein family, such as eosin Y, a tetrabrominated fluorescein, provided similar results as the parent compound fluorescein (14% yield, 97:3 e.r., entry 5). Interestingly, a dibrominated analog of fluorescein, dibromofluorescein, demonstrated superior activity compared to both fluorescein and eosin Y, providing azidation product **2a** in 23% yield while maintaining excellent enantioselectivity (98:2 e.r., entry 6). Reducing the loading of dibromofluorescein to 1 mol% slowed down photoinduced free radical generation and resulted in a further improved yield of 36% (entry 7). It was found that only 1 to 2 equiv of NaN_3_ was required for this tyrosinase-catalyzed asymmetric radical azidation (Supplementary Table 6), indicating enhanced azide binding affinity of binuclear copper tyrosinases relative to nonheme Fe enzymes^19,20^. Importantly, lowering the NaN_3_ increased the yield of **2a** to 40% by suppressing the non-enzymatic formation of acylazide side products derived from **1a** (entry 8). Irradiation at 520 nm further improved the yield of **2a**, due to the stronger absorption of dibromofluorescein at this wavelength (entries 9 and 10). Carrying out this tyrosinase-catalyzed azidation in the presence of added transition-metal ions including Mn^2+^, Fe^2+^, Co^2+^ and Ni^2+^ spanning a wide range of concentrations of did not lead to azidation product formation (entry 13), confirming the critical role of copper in this unnatural radical transformation.

### Protein engineering and substrate scope of binuclear Cu tyrosinase-catalyzed enantioselective radical azidation

With the optimized conditions established for wild-type *Bm*Tyr, we next performed semi-rational protein engineering by targeting active-site and substrate entrance tunnel hydrophobic residues, with the aim to modify the active site to better accommodate the hydrophobic alkyl radical intermediates in the new-to-nature chemistry. Guided by the crystal structure of *Bm*Tyr (PDB ID: 6EI4)^56^, residues proximal to the binuclear Cu centers supported by six histidine residues (H42, H60 and H69 for Cu1; H204, H208 and H231 for Cu2) were targeted to construct a mutant library to investigate the effects of active-site residues on non-native azidation. It was found that the inclusion of a valine to alanine mutation at residue 218, residing in a flexible loop flanking the binuclear Cu centers (V218A), was beneficial, leading to an improved azidation yield of 56% (entry 2). By reducing the size of the amino acid residue 218 close to the Cu centers, V218A better accommodated organic substrates to approach the reactive Cu centers. Using *Bm*Tyr V218A as the parent, another beneficial mutation N205M adjacent to the Cu-binding residue H204 was discovered, further enhancing the yield of **2a** to 70% (entry 3). Collectively, the engineered tyrosinase double mutant *Bm*Tyr V218A N205M allowed this non-native photometallobiocatalytic radical azidation to occur with improved efficiency in an enantioselective fashion. Importantly, ablating either Cu center from the binuclear Cu enzyme by mutating all three Cu-binding histidine residues to alanine (*Bm*Tyr V218A N205M H42A H60A H69A, entry 4; *Bm*Tyr V218A N205M H204A H208A H231A, entry 5) led to drastically diminished activity and enantioselectivity, highlighting the essential role of the binuclear Cu center in facilitating this asymmetric radical azidation.

With this engineered *Bm*Tyr V218A N205M double mutant in hand, we next examined the substrate scope of radical decarboxylative azidation (Fig. 3b). It was found that redox-active esters bearing a *para*- (**2b**), a *meta*- (**2c**) and an *ortho*- (**2d**) substituent on the aromatic ring were compatible with the engineered binuclear Cu biocatalyst. Halogen substituents including a fluoro (**2e**), a chloro (**2f**), and a bromo (**2g**) were readily accommodated under these photobiocatalytic conditions. Electron-donating groups such as a methoxy (**2h**) and electron-withdrawing groups such as a trifluoromethyl (**2i**) at the *para*-position of the aromatic ring were also compatible, although reduced enantioselectivity was observed for the *para*-methoxy substrate. Additionally, our engineered *Bm*Tyr V218A N205M variant could also readily transform an extended α-ethyl substrate (**2j**), indicating the potential of this binuclear Cu enzyme to accommodate larger substrates.

**Fig. 3.**
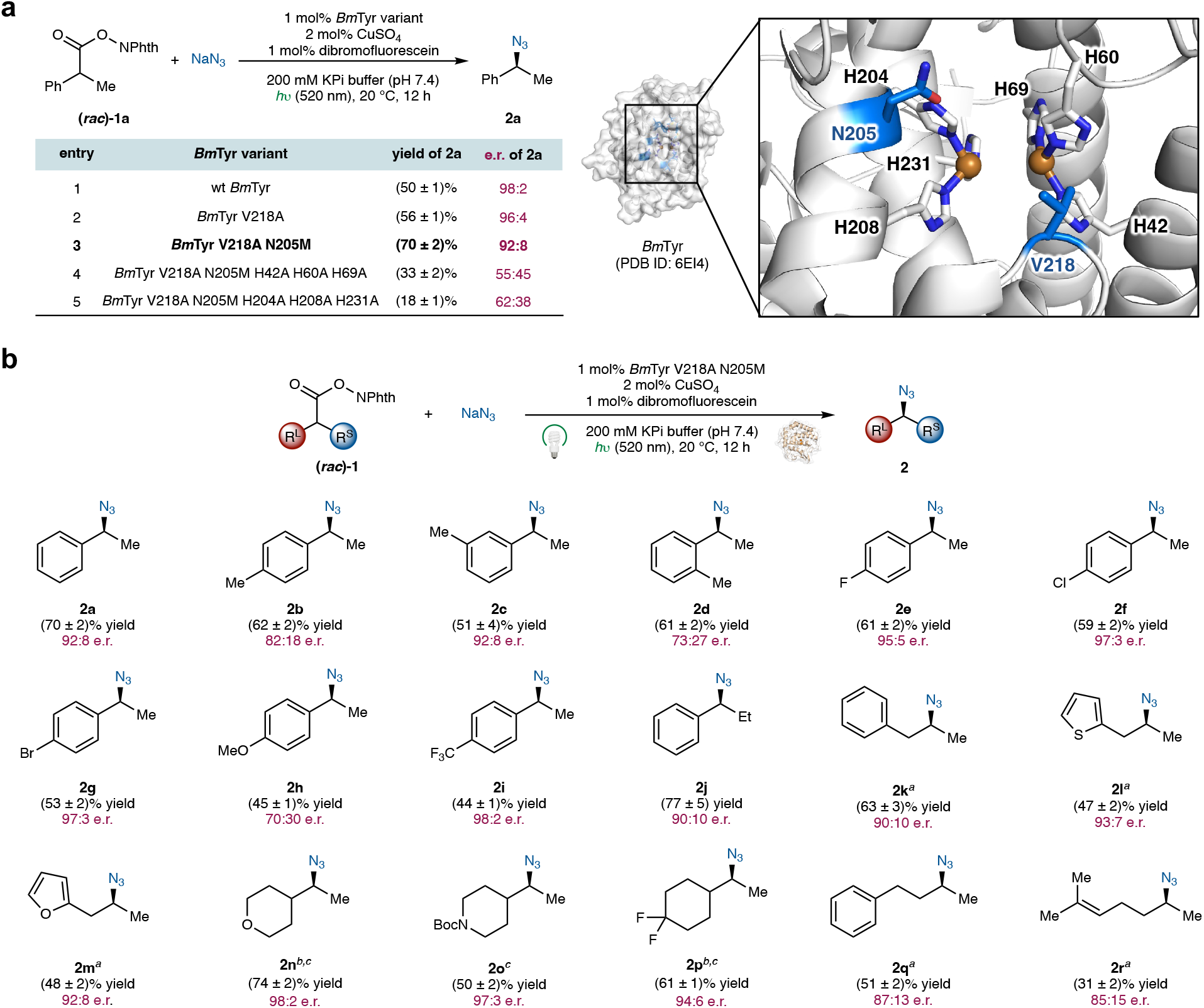
Engineering of the binuclear Cu tyrosinase from *Bacillus megaterium* for enantioselective photometallobiocatalytic radical azidation. **a**, Protein engineering of *Bm*Tyr for the enantioselective decarboxylative azidation. The active-site illustration of *Bm*Tyr was generated from 6EI4 (PDB ID) using PyMOL. Mutated residues are highlighted in marine. **b**, Substrate scope of enantioselective photometallobiocatalytic decarboxylative azidation. Reaction conditions: *N*-hydroxyphthalimide ester **1** (1.0 equiv, 4.0 mM), NaN_3_ (2 equiv, 8.0 mM), 1 mol% *Bm*Tyr V218A N205M (40 μM), 2 mol% CuSO_4_ (80 μM), 1 mol% dibromofluorescein (40 μM), *h*υ (520 nm), KPi buffer (200 mM, pH 7.4), 20 °C, 12 h. Biocatalytic reactions were carried out in triplicate and average yields and standard deviations were reported. See the Supplementary Information for details. ^*a*^1 mol% wt *Bm*Tyr was used. ^*b*^1 mol% *Bm*Tyr R209M was used. ^*c*^440 nm in lieu of 520 nm.

In cooperative photobiocatalysis, the efficient and enantioselective conversion of unstabilized alkyl radicals, namely radicals lacking a neighboring stabilizing group such as an aryl and an α-carbonyl substituent, continues to present a significant challenge due to their short lifetimes. In particular, with previously developed nonheme Fe biocatalyst systems, transformations involving these unstabilized aliphatic radicals did not provide synthetically useful yields or good stereo-control^19^. We therefore investigated whether binuclear copper enzymes could enable the conversion of these reactive radical intermediates in a productive manner. It was found that wild-type *Bm*Tyr allowed the transformation of homobenzylic substrates to the corresponding azidation products (**2k–2m**) in good yields and enantioselectivities. Heterocycles, including a thiophene (**2l**) and a furan (**2m**), were compatible under these reaction conditions. Furthermore, fully aliphatic substrates lacking an adjacent stabilizing aromatic substituent, such as aliphatic substrates bearing a tetrahydropyran (**2n**), a piperidine (**2o**) and a *gem*-difluorinated cyclohexyl (**2p**) moiety, could also be effectively transformed by *Bm*Tyr R209M or *Bm*Tyr V218A N205M with excellent enantioselectivity. Additional secondary alkyl substrates bearing a phenylethyl (**2q**), and an isopentenyl (**2r**) group at the α carbon, could also be transformed in an enantioselective fashion. Importantly, the ability to effectively distinguish between two similar alkyl substituents represented an ongoing challenge in asymmetric alkyl cross-coupling reactions^46,47^, and our results highlighted the excellent potential of binuclear copper enzymes to solve difficult problems in asymmetric catalysis. Furthermore, the efficient and enantioselective conversion of unstabilized alkyl radicals with binuclear copper enzymes is highly complementary to nonheme Fe-based enzymatic systems.

### Tyrosinase-catalyzed enantioselective radical bromination

Compared to azidation, biocatalytic radical bromination via an unnatural mechanism remains challenging, in part due to the substantially lower binding affinity of the bromide ion for metallocofactors in aqueous solution. Furthermore, catalytic asymmetric radical bromination with small-molecule transition-metal catalysts applicable to a broad range of organic substrates has remained a difficult task in enantioselective catalysis. After establishing the tyrosinase-catalyzed enantioselective azidation, we next questioned whether this radical rebound reactivity could be expanded to bromides, giving rise to unnatural copper-dependent halogenases, an activity which is exceedingly rare among natural copper enzymes. Previous studies showed that halide ions are potent inhibitors for tyrosinases in their native activity due to the bridging μ_2_-binding mode of the halide anion with both copper centers^42,43^. Although this strong binding is deleterious to the native tyrosinase activity, we posited that such enhanced bromide binding with a binuclear Cu–Br–Cu intermediate might enable otherwise highly challenging radical bromination reactions. Indeed, using (*rac*)-**1n** as the model substrate, in the presence of 40 equiv bromide ion, the use of 1 mol% dibromofluorescein and 1 mol% wt *Bacillus megaterium* tyrosinase provided the desired alkyl bromide product **3a** in 47% yield and 88:12 e.r. (Fig. 4a, entry 1). Re-evaluation of a wider range of photosensitizers revealed that the parent unsubstituted fluorescein provided improved yield and enantioselectivity (64% yield and 90:10 e.r., entry 2). To further enhance the activity of this dicopper brominase, we evaluated a focused collection of *Bm*Tyr variants. It was found that the R209M mutant of *Bm*Tyr furnished further improved activity, giving rise to **3a** in 80% yield and 90:10 e.r. (entry 3). Active-site residue 209 lies close to the copper-binding H208, and the inclusion of R209M may facilitate copper recruitment and binding in the dicopper binding site^60^. Similar to our observation for biocatalytic radical azidation, eliminating either Cu center from the binuclear Cu system (*Bm*Tyr R209M H42A H60A H69A, entry 4; *Bm*Tyr R209M H204A H208A H231A, entry 5) drastically reduced the activity and enantioselectivity of the present Cu-dependent biocatalyst.

**Fig. 4.**
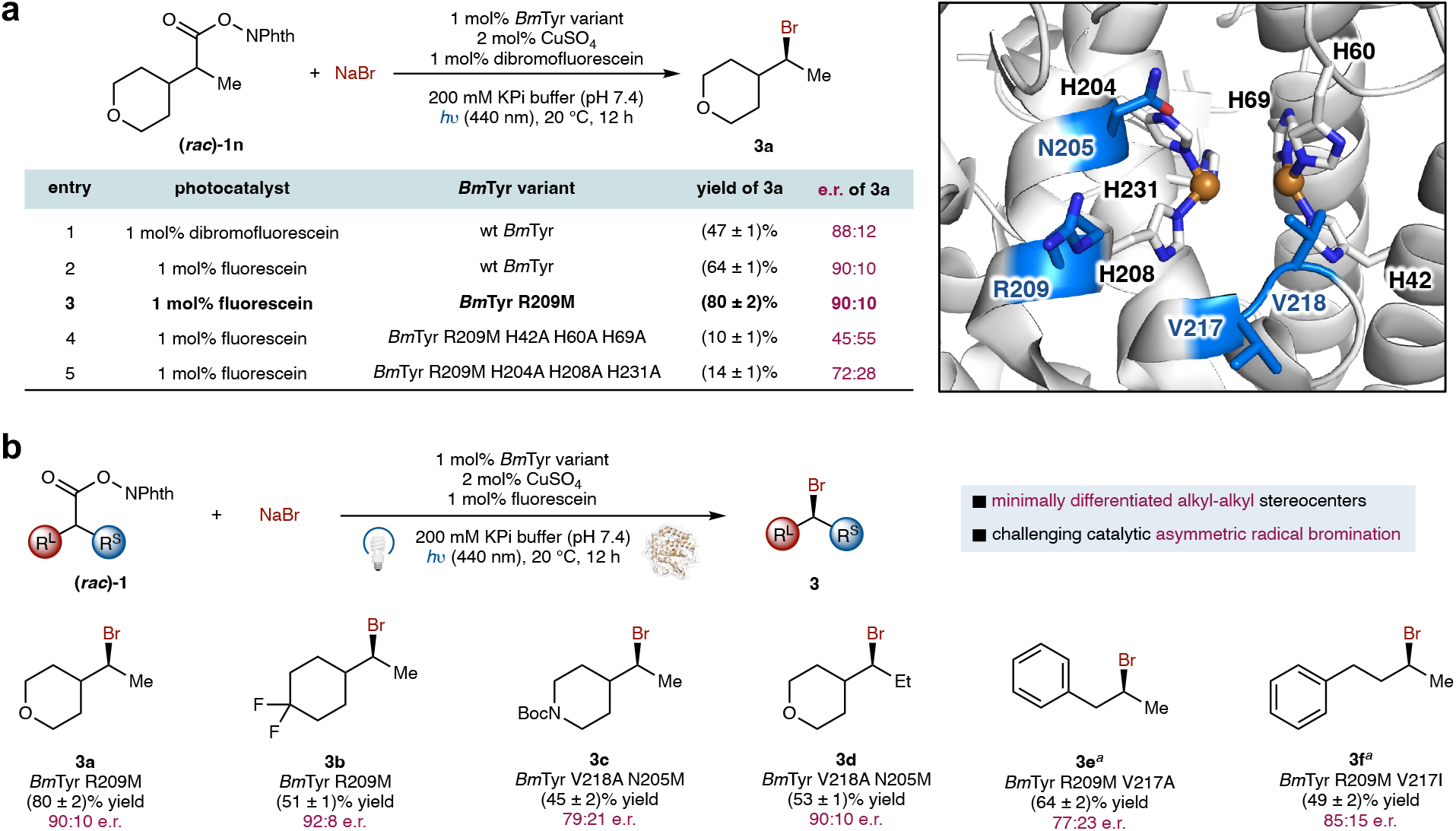
Binuclear copper enzyme-catalyzed enantioselective decarboxylative bromination of redox-active esters. **a**, Optimization of photobiocatalytic asymmetric decarboxylative bromination of redox-active esters. The active-site illustration of *Bm*Tyr was generated from 6EI4 (PDB ID) using PyMOL. Mutated residues were highlighted in marine. **b**, Substrate scope of photobiocatalytic asymmetric decarboxylative bromination of redox-active esters. Reaction conditions: redox-active ester **1** (1.0 equiv, 4.0 mM), NaBr (40 equiv, 160 mM), 1 mol% *Bm*Tyr variant (40 μM), 2 mol% CuSO_4_ (80 μM), 1 mol% fluorescein (40 μM), *h*υ (440 or 520 nm), 200 mM KPi buffer (pH 7.4), 20 °C, 12 h. Biocatalytic reactions were carried out in triplicate and average yields and standard deviations were reported. See the Supplementary Information for details. ^*a*^520 nm in lieu of 440 nm.

With these optimized reaction conditions in hand, we surveyed the substrate scope of this enantioselective photobiocatalytic decarboxylative bromination. It was found that functionalized cyclic alkyl substituents were compatible with this transformation. For example, a substrate bearing a 4,4-*gem*-difluorocyclohexyl group (**3b**) could be transformed in excellent enantioselectivity (92:8 e.r.). *N*-Boc-piperidine (**3c**) and tetrahydropyran (**3d**) substituents were also readily accommodated. Using *Bm*Tyr V218A N205M possessing a further expanded active-site, the more challenging enantiodifferentiation between a tetrahydropyran group and an ethyl group (**3d**) could also be realized with good levels of enantiocontrol (90:10 e.r.). Furthermore, without additional protein engineering, homobenzylic (**3e**) and 2-(4-phenylbutyl) (**3f**) substrates also underwent this photobiocatalytic transformation with promising levels of enantioselectivity. Given the high similarity between the methyl and the primary alkyl substituent flanking the radical center, these results further underscored the excellent potential of binuclear copper enzymes in tackling challenging enantioinduction problems. Further enzyme engineering to improve enantioselectivity through high-throughput experimentation is currently underway in our laboratory.

### Tyrosinase-catalyzed enantioselective radical chlorination and isothiocyanation

Furthermore, we found that binuclear copper enzymes could also facilitate other types of radical halogenation reactions, including chlorination and isothiocyanation (Fig. 5). Using racemic redox-active ester (*rac*)-**1n** as the substrate, in the presence of 40 equiv chloride ion, wt *Bm*Tyr catalyzed enantioselective radical chlorination to afford the corresponding alkyl chloride in 53% yield and 85:15 e.r. (Fig. 5a). When sodium thiocyanate was used, the binuclear copper enzyme *Bm*Tyr transformed (*rac*)-**1n** into the isothiocyanated product (**5**, 37% yield, 94:6 e.r.) as the major product, with an isothiocyanation : thiocyanation ratio of 83:17 (Fig. 5b). SCN− is an ambident anion, and literature studies are consistent with nitrogen binding to copper in small-molecule systems^61^. Importantly, this binuclear Cu enzyme furnished an ambident selectivity complementary to nonheme Fe enzyme–catalyzed radical transformations using thiocyanate anion, where our engineered *Pseudomonas putida* metapyrocatechase (*Pp*MPC) variant *Pp*MPC I291L L155F I204L afforded the alkyl thiocyanate product exclusively in 54% yield and 95:5 e.r.^20^. Collectively, these results revealed the distinct reactivity of copper enzymes in radical functionalization reactions, highlighting the complementary activity profiles of Cu and Fe-based nonheme enzyme systems.

**Fig. 5.**
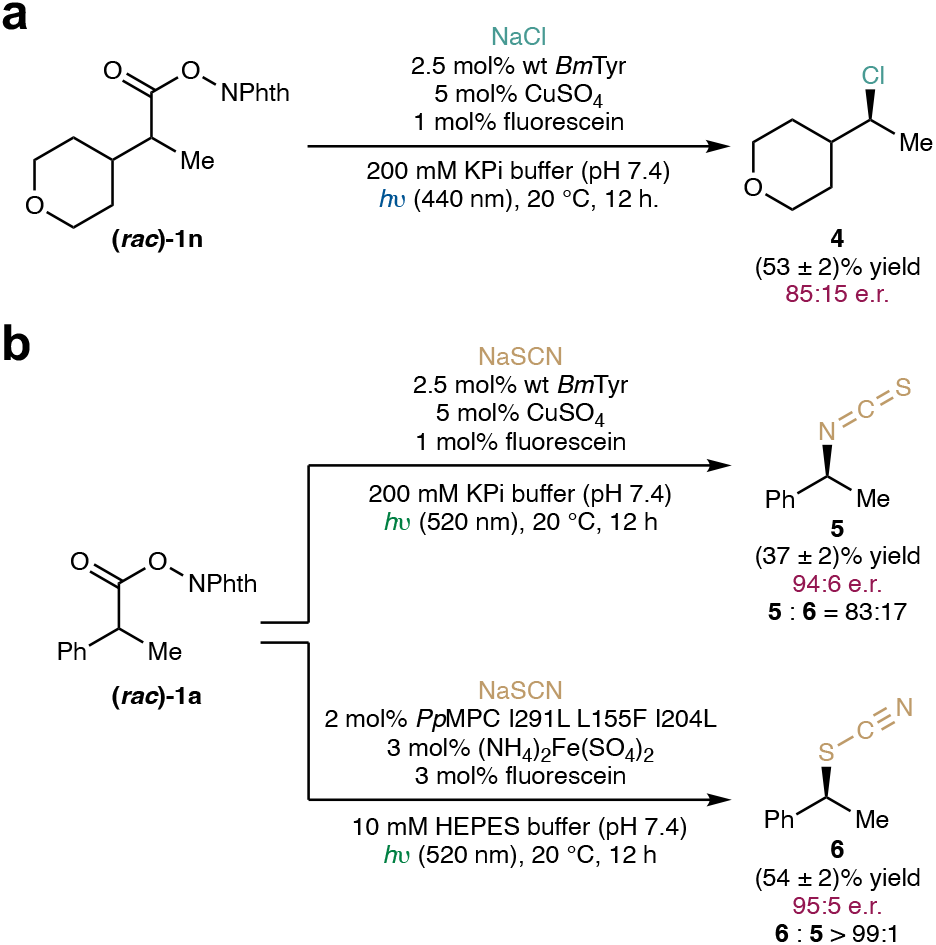
Binuclear copper enzyme-catalyzed enantioselective decarboxylative chlorination and isothiocyanation. Chlorination conditions: redox-active ester **1n** (1.0 equiv, 4.0 mM), NaCl (40.0 equiv, 160 mM), 2.5 mol% wt *Bm*Tyr (100 μM), 5 mol% CuSO_4_ (200 μM), 1 mol% fluorescein (40 μM), *h*υ (440 nm), 200 mM KPi buffer (pH 7.4), 20 °C, 12 h. Isothiocyanation conditions: redox-active ester **1a** (1.0 equiv, 4.0 mM), NaSCN (5.0 equiv, 20 mM), 2.5 mol% wt *Bm*Tyr (100 μM), 5 mol% CuSO_4_ (200 μM), 1 mol% fluorescein (40 μM), *h*υ (520 nm), 200 mM KPi buffer (pH 7.4), 20 °C, 12 h.

### EPR and UV–visible spectroscopic characterization of dinuclear Cu enzymes

Tyrosinases typically feature a coupled binuclear dicopper active site, in which each copper ion is coordinated by three conserved histidine residues^28,62^. However, *Bm*Tyr displays distinct copper-binding behavior. Its crystal structure reveals the presence of both dicopper and monocopper active sites^63^. Consistent with this observation, titration of two equivalents of CuSO_4_ followed by removal of unbound copper using a desalting column resulted in clear mononuclear copper EPR signals in both wild-type and mutant *Bm*Tyr (Figs. 6a and 6b). EPR spectral simulations indicate that both variants share a similar mononuclear copper site, characterized by *g* values of (2.24, 2.07, 2.05) and a copper hyperfine coupling of 190 × 10−^4^ cm−^1^ (Figs. 6a and 6b), consistent with a type 2 copper center with d_x2-y2_ ground state^28^.

**Figure 6.**
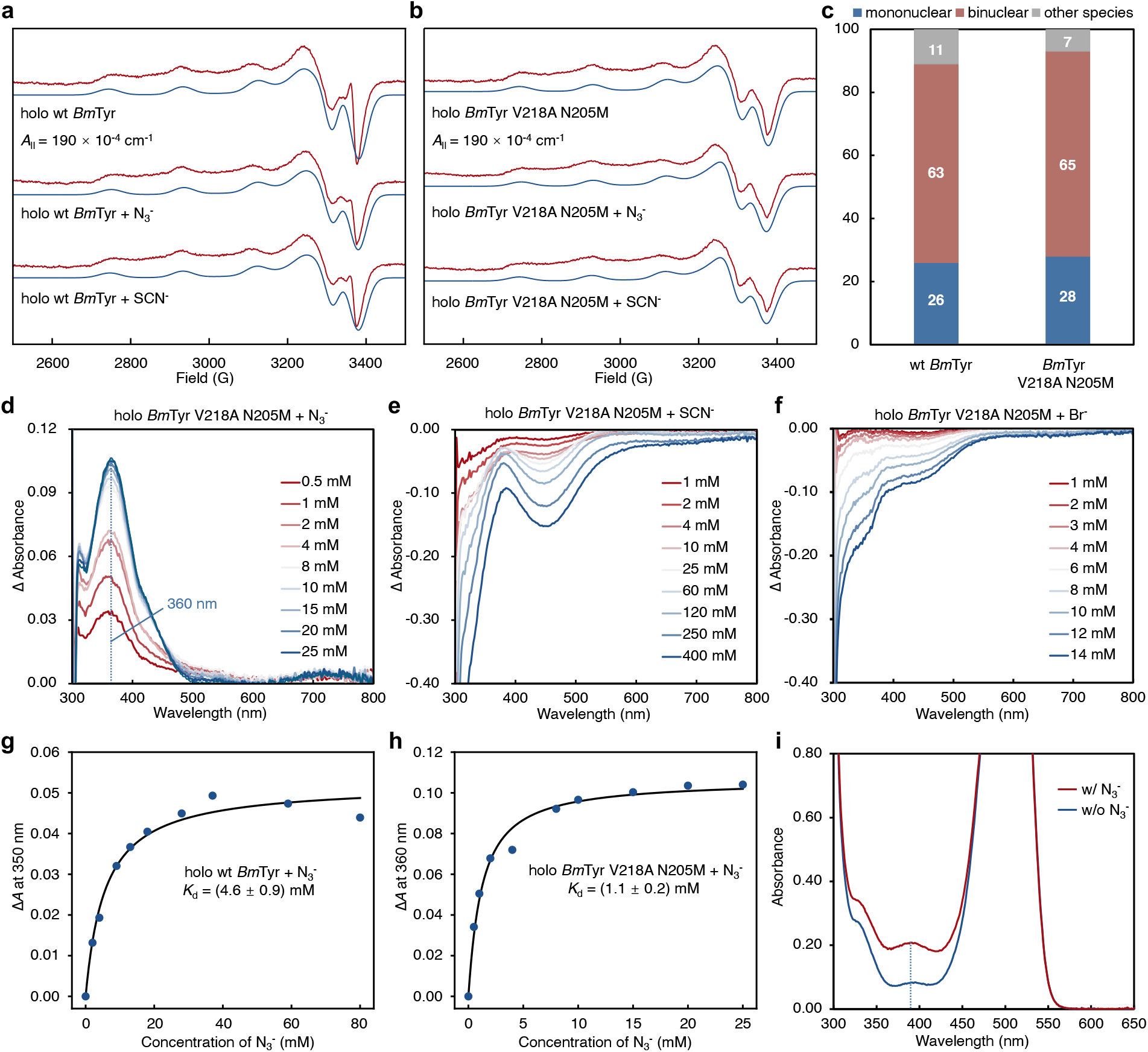
Spectroscopic characterization of copper speciation and (pseudo)halide binding in wild-type and mutant *Bm* tyrosinase. **a**, Experimental (red) and simulated (blue) X-band EPR spectra of wt *Bm*Tyr in the absence and presence of ligands (N_3_− and SCN−). **b**, Experimental (red) and simulated (blue) X-band EPR spectra of *Bm*Tyr V218A N205M in the absence and presence of ligands (N_3_− and SCN−). **c**, Distribution of copper species in wt *Bm*Tyr and *Bm*Tyr V218A N205M determined by spin-counting EPR and biquinoline assay, showing the average percentages of mononuclear and binuclear copper species from triplicate measurements. Error bars represent standard deviations (wt *Bm*Tyr: ±5.2% mononuclear, ±0.6% binuclear; *Bm*Tyr V218A N205M: ±8.3% mononuclear, ±4.6% binuclear). **d**, UV–visible difference spectra (*ΔA*) from azide titration of *Bm*Tyr V218A N205M in 50 mM Tris buffer, referenced to the 0 mM azide spectrum. **e**, UV– visible difference spectra (*ΔA*) from thiocyanate titration of *Bm*Tyr V218A N205M in 50 mM Tris buffer, referenced to the 0 mM thiocyanate spectrum. **f**, UV–visible difference spectra (*ΔA*) from bromide titration of *Bm*Tyr V218A N205M in 50 mM Tris buffer, referenced to the 0 mM bromide spectrum. **g**, Azide binding isotherm for wt *Bm*Tyr derived from absorbance changes at 350 nm. **h**, Azide binding isotherm for *Bm*Tyr V218A N205M derived from absorbance changes at 360 nm. **i**, *In situ* UV–visible spectroscopy of photobiocatalytic decarboxylative azidation with *Bm*Tyr V218A N205M.

To quantify the distribution between mono- and dicopper species under catalytic conditions, we combined spin-counting EPR with a colorimetric biquinoline assay. The coupled binuclear type III copper site is EPR-silent in its Cu(II)_2_ state due to strong antiferromagnetic coupling gives an overall *S* = 0 ground state, whereas mononuclear copper remains EPR-active in its Cu(II) form. Thus, spin-counting EPR was used to determine the concentration of mononuclear copper, while the biquinoline assay provided the total copper concentration. The amount of coupled binuclear copper was then calculated as the difference between total and mononuclear copper. As shown in Fig. 6c, the total copper concentration in the *Bm*Tyr V218A N205M is 291 μM, of which 42 μM corresponds to mononuclear copper. This yields a coupled binuclear copper concentration of 124 μM (two Cu per site). These results indicate that, under catalytic conditions, 65% of *Bm*Tyr V218A N205M is populated by dicopper sites, whereas 28% contains a single copper ion. A similar distribution is observed in the wild-type *Bm*Tyr, with 63% and 26% of the enzyme populated by di- and mononuclear copper sites, respectively (Fig. 6c). Additionally, it was found that *holo* enzyme reconstitution with Cu(MeCN)_4_BF_4_ under anaerobic conditions led to nearly quantitative formation of binuclear Cu(I) species, as determined by biquinoline assays. The reconstituted, and air-treated *Bm*Tyr enzyme sample displayed similar catalytic activity and enantioselectivity in photobiocatalytic azidation and bromination (see Supplementary Figs. 15 and 16 for details).

Although the coupled binuclear copper site is the predominant form of mutant *Bm*Tyr under catalytic conditions, we next sought to determine which copper species—binuclear or mononuclear—is catalytically relevant. Because the mononuclear copper site is EPR-active, we first examined (pseudo)halide binding to both mutant and wild-type enzymes using EPR spectroscopy. As shown in Fig. 6a, addition of a 20-fold excess of azide or thiocyanate resulted in no detectable perturbation in the EPR spectra, suggesting that the mononuclear copper site either does not bind (pseudo)halides or binds them only very weakly. Thus, the mononuclear copper species is likely not involved in catalysis under our photobiocatalytic conditions. We then probed (pseudo)halide binding to the coupled binuclear copper site using UV–visible absorption spectroscopy. Upon addition of azide to the mutant enzyme, a new ligand-to-metal charge-transfer (LMCT) band at 360 nm was observed (Fig. 6d, see Supplementary Figs. 8 and 10 for details). Similar LMCT features were reported for azide binding in both tyrosinases and synthetic dicopper complexes^41,64-66^. Fitting of the titration data provided a dissociation constant (*K*_d_) of (4.6 ± 0.9) mM for wt *Bm*Tyr and (1.1 ± 0.2) mM for *Bm*Tyr V218A N205M respectively (Figs. 6g and 6h). This value is consistent with reported affinities for azide binding to binuclear copper sites in tyrosinases (typically 1–2 mM)^67^. In contrast, azide binding to mononuclear copper centers is significantly weaker; for example, reported *K*_d_ values are 11 mM for Cu/Zn superoxide dismutase^68^ and 18 mM for the E2 domain of the amyloid precursor protein^69^. Consistent with this assignment, titration with SCN− and Br− also induced perturbations in the LMCT at 330 nm characteristic of the type-3 copper site in the mutant enzyme (Figs. 6e and 6f, see Supplementary Figs. 12 and 14 for details). Although the lower levels of binding affinities for these ligands preclude the determination of dissociation constants under our conditions, their spectral effects support analogous interaction with the binuclear copper center. This behavior is also consistent with previous studies showing that halide ions inhibit tyrosinase activity through binding at the dicopper active site^42^. Furthermore, *in situ* UV–visible spectroscopy was used to probe the formation of binuclear Cu azide intermediates under catalytically relevant conditions in the presence of the redox-active ester substrate, dibromofluorescein and NaN_3_. During the product-forming, catalytic turnover conditions, an absorption at 360 nm corresponding to the binuclear Cu azide charge transfer band was observed. This feature was not present before the addition of NaN_3_ (Fig. 6i, see Supplementary Figs. 19 and 20 for details). These results further confirmed the formation of the binuclear Cu azide intermediate under photobiocatalytic conditions, further supporting its involvement in catalysis. Taken together, these results establish that the coupled binuclear copper site is both the dominant species under catalytic conditions and the catalytically relevant center for the new-to-nature (pseudo)halogenation chemistry, enabling (pseudo)halide binding and transfer to the photoredox-generated radical intermediate.

Additionally, to probe the origin of activity difference between structurally related photosensitizers fluorescein, dibromofluorescein and Eosin Y. We performed fluorescence anisotropy titration to determine the binding affinity between the photocatalysts and *Bm*Tyr V218A N205M used in our study (Extended Data Fig. 1). Fluorescence anisotropy titration with fluorescein showed no saturation within the investigated concentration regime, indicating an estimated *K*_d_ of (850 ± 50) μM. In contrast, the *K*_d_ value of dibromofluorescein with *Bm*Tyr V218A N205M was determined to be (14.0 ± 1.0) μM. Furthermore, Eosin Y, a tetrabrominated fluorescein derivative, exhibited a further enhanced binding affinity, with a *K*_d_ of (5.1 ± 0.2) μM. These results indicated a substantially stronger binding of halogenated fluorescein congeners, including dibromofluorescein and Eosin Y, relative to that of fluorescein. Thus, we ascribe the contrasting activity of these structurally related dyes in part to their differential binding affinities toward the metalloenzyme catalyst.

## Conclusion

In summary, we established binuclear copper enzymes as a powerful platform for enabling new-to-nature stereoselective radical transformations. Through the investigation of diverse natural copper enzymes, we demonstrated that type III dicopper tyrosinases could be repurposed to catalyze enantioselective radical (pseudo)halogenation reactions, including bromination, chlorination, azidation and isothiocyanation. Mechanistic and spectroscopic studies revealed that the binuclear copper center plays a central role in facilitating these transformations, providing both efficient (pseudo)halide binding and a favorable environment for radical coupling. These findings highlight the unique reactivity of bimetallic enzymes enabled by multimetallic cooperativity in the context of new-to-nature enzymology, opening up new avenues for advancing asymmetric radical transformations that remain challenging for small-molecule catalysts and mononuclear metalloenzymes. In particular, the ability of dicopper enzymes to engage and transform highly reactive, unstabilized alkyl radicals into products bearing a minimally differentiated alkyl-alkyl stereocenter addresses a longstanding challenge in asymmetric catalysis. We anticipate that the integration of multimetallic enzyme scaffolds with non-native reaction design will unlock previously inaccessible transformations at the interface of biocatalysis and synthetic chemistry.

## Methods

### Expression of *Bm*Tyr variants

*E. coli* BL21(DE3) cells harboring recombinant plasmid encoding the appropriate *Bm*Tyr variant were grown in LBamp media (50 mL, with 0.10 mg/mL ampicillin) at 37 °C and 250 rpm for ca. 6 h, until OD_600_ reached 0.6–0.8 (mid to late log phase). Preculture (50.0 mL, 5% v/v) was used to inoculate 1 L TB media supplemented with 0.10 mg/mL ampicillin (TB_amp_) in a 2 L Erlenmeyer flask. The culture was incubated at 37 °C and 230 rpm for 3 h to reach an OD_600_ of ca. 1.5. The culture was then cooled on ice for 20 min and induced with 0.5 mM isopropyl β-D-1-thiogalactopyranoside (IPTG, final concentrations). Protein expression was performed at 30 °C and 200 rpm for 16 h. The cells were then transferred to a conical tube (50 mL) and harvested by centrifugation (3,434 *g*, 4 min, 4 °C) using an Eppendorf 5910R tabletop centrifuge.

### General Procedure for enantioselective photobiocatalytic azidation

Purified *Bm*Tyr variant (ca. 29 mg/mL, 0.8 mM, 50 μL) was allowed to thaw and kept on ice. In a Coy anaerobic chamber, the following stock solutions were prepared: redox-active ester **1** (200 mM in degassed DMSO), dibromofluorescein (2.0 mM in degassed DMSO), NaN_3_ (400 mM in degassed dd H_2_O), CuSO_4_ (4.0 mM in degassed dd H_2_O). To a one-dram vial containing a stir bar were added 890 μL degassed KPi buffer (200 mM, pH 7.4), ca. 50 μL *Bm*Tyr variant (the volume of the *Bm*Tyr stock solution was adjusted according to the concentration of the enzyme sample) and 20 μL CuSO_4_ stock solution, which was allowed to incubate for 1 min on a microplate shaker at 600 rpm. Followed by the addition of 20 μL redox-active ester **1** stock solution, 20 μL NaN_3_ stock solution and 20 μL dibromofluorescein stock solution. The final reaction volume was 1000 μL; the final concentrations of each reaction component were as follows: 4.0 mM of redox-active ester **1**, 8.0 mM NaN_3_, 80 μM CuSO_4_, 40 μM *Bm*Tyr variant and 40 μM dibromofluorescein. The vials were sealed, removed from the Coy anaerobic chamber and placed in a Shanghai 3S photoreactor (300 rpm, 20 °C) and illuminated at the indicated wavelength (440 nm or 520 nm). After 12 h, 1000 μL of a mesitylene solution (0.50 mM in EtOAc, used for chiral GC analysis) or (0.50 mM in *i*-Pr_2_O, used for chiral HPLC analysis) was added to the mixture. The vial was sealed with a screw cap and vortexed. The resulting mixture was transferred to a 2 mL microcentrifuge tube and centrifuged (15,000 rpm, 10 min) using an Eppendorf 5424R centrifuge to separate the organic and aqueous layers. The organic layer (600 μL) was transferred to a 2 mL GC vial and analyzed by chiral GC or HPLC.

### General Procedure for enantioselective photobiocatalytic bromination

Purified enzyme (ca. 29 mg/mL, 0.8 mM, 50 μL) was allowed to thaw and kept on ice. In a Coy anaerobic chamber, the following stock solutions were prepared: redox-active ester **1** (200 mM in degassed DMSO), fluorescein (2.0 mM in degassed DMSO), NaBr (178 mM in 200 mM KPi buffer, pH 7.4), CuSO_4_ (4.0 mM in degassed dd H_2_O). To a one-dram vial containing a stir bar were added 900 μL NaBr stock solution, ca. 50 μL *Bm*Tyr variant (the volume of the *Bm*Tyr stock solution was adjusted according to the concentration of the enzyme sample) and 20 μL CuSO_4_ stock solution, which was allowed to incubate for 1 min on a microplate shaker at 600 rpm. Followed by the addition of 20 μL redox-active ester **1** stock solution and 20 μL fluorescein stock solution. The final reaction volume was 1000 μL; the final concentrations of each reaction component were as follows: 4.0 mM redox-active ester **1**, 160 mM NaBr, 80 μM CuSO_4_, 40 μM *Bm*Tyr variant and 40 μM fluorescein. The vials were sealed, removed from the Coy anaerobic chamber and placed in a Shanghai 3S photoreactor (300 rpm, 20 °C) and illuminated with indicated wavelength (440 nm or 520 nm). After 12 h, 1000 μL of a mesitylene solution (0.50 mM in EtOAc, used for chiral GC analysis) or (0.50 mM in *i*-Pr_2_O, used for chiral HPLC analysis) was added to the mixture. The vial was sealed with a screw cap and vortexed. The resulting mixture was transferred to a 2 mL microcentrifuge tube and centrifuged (15,000 rpm, 10 min) using an Eppendorf 5424R centrifuge to separate the organic and aqueous layers. The organic layer (600 μL) was transferred to a 2 mL GC vial and analyzed by chiral GC or HPLC.

## Acknowledgements

We are grateful to the National Institutes of Health (R35GM147387 to Y.Y. and R35GM155016 to S.T.) for supporting the experimental work and the National Science Foundation (CHE-2400087 to P.L.) for supporting the computational work. Y.Y. is an Alfred P. Sloan Research Fellow (FG-2024-22244), a Camille Dreyfus Teacher-Scholar Awardee (TC-25-084), a David & Lucile Packard Fellow (2023-76169) and a Howard Hughes Medical Institute Freeman Hrabowski Scholar. Computations were carried out at the University of Pittsburgh Center for Research Computing and Data and the Advanced Cyberinfrastructure Coordination Eco-system: Services & Support (ACCESS) Program, supported by NSF awards OAC-2117681, OAC-1928147, OAC-1928224 and CHE-260005.

## Data availability

All data are available in the main text and the Supplementary Information. Plasmids encoding engineered tyrosinase variants reported in this study are available for research purposes from Y.Y. under a material transfer agreement with the University of California Santa Barbara.

## Author contributions

Y.Y. conceived and directed the project. X.-W.C., H.-Y.C., and Y.Y. performed Cu enzyme mining, cloning and overexpression in *E. coli*. X.-W.C. performed Cu enzyme evaluation, engineering, substrate scope studies and substrate and racemic product synthesis. H.-D.C., L.-P.Z., and K. L. prepared additional substrates and racemic products. B.K.M. carried out the computational studies with P.L. providing guidance. O.M.A., N.N., J.P., and S.T. carried out spectroscopic and biochemical characterization of Cu enzymes. X.-W.C., O.M.A., Y.Y., and S.T. wrote the manuscript with the input of all other authors.

## Competing interests

Y.Y. and X.-W.C. are inventors on a patent application submitted by the University of California Santa Barbara that covers compositions, systems, and methods for biocatalytic (pseudo)halogenation with Cu enzymes. The remaining authors declare no competing interests.

---

**Extended Data Fig 1.**
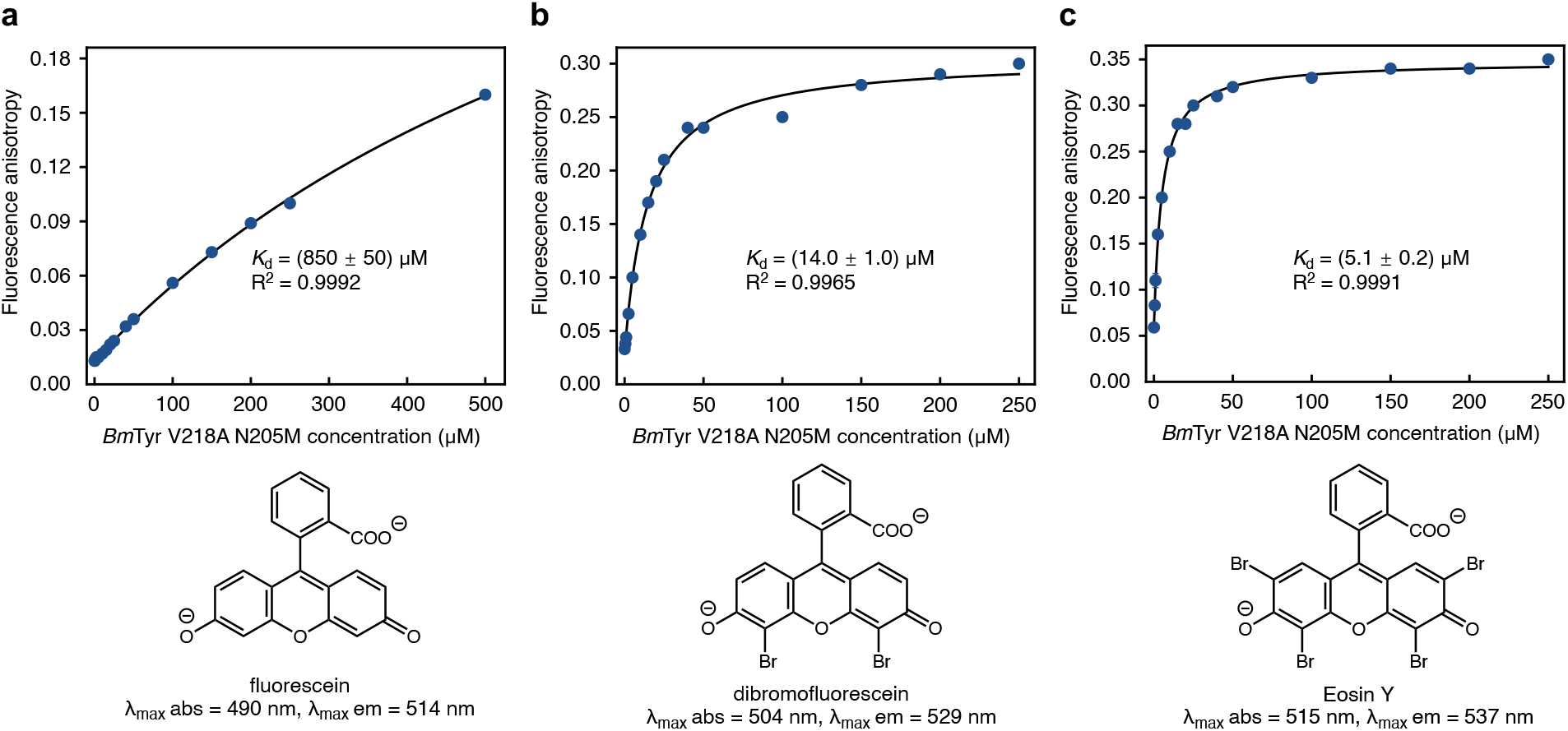
Fluorescence anisotropy titration measurements with *Bm*Tyr V218A N205M. **a**, Determination of *K*_d_ for fluorescein binding to *Bm*Tyr V218A N205M. **b**, Determination of *K*_d_ for dibromofluorescein binding to *Bm*Tyr V218A N205M. **c**, Determination of *K*_d_ for Eosin Y binding to *Bm*Tyr V218A N205M.

## Notes

### Competing Interest Statement

The authors have declared no competing interest.

